# High-resolution Cell Atlas of Domestic Pig Lung and Online Platform Exploring for Lung Single Cell

**DOI:** 10.1101/2020.12.20.423716

**Authors:** Lijing Zhang, Jiacheng Zhu, Haoyu Wang, Jun Xia, Ping Liu, Fang Chen, Hui Jiang, Weiying Wu, Lihua Luo, Dongsheng Chen, Xingliang Zhang

## Abstract

Due to the comparable organ sizes and physiology to human beings, genetically engineered pig is gradually regarded as an optimal source of human for transplantation and an excellent model for human diseases research. The major barrier between pig and human is mainly cellular heterogeneity and immune incompatibilities. Myriad scRNA-seq data of human has been reported, but the counterpart in pig is scarce. Here, we applied scRNA-seq technology to study the cellular heterogeneity of 3 months old pig lungs, generating the single-cell atlas of 13,580 cells covering 16 major cell types. Based on this data, we systematically characterized transcription factor regulatory networks, cell-cell communications in each cell type, and comparison of them and those in human lung. We first presented a comprehensive and openly accessible online platform, ScdbLung, which was composed of three functional modules (Marker, Cluster, and Download) and dedicated to exploiting the valuable resources of lung scRNA-seq data across four mammalian species. Overall, our scRNA-seq atlas of domestic pig lung, conserved transcription factor regulatory networks and cell-cell communications between domestic pig and human, and ScdbLung could provide guidance for porcine lung research and clinical applicability.

## Introduction

The mammalian lung is a complex tissue with high cellular heterogeneity, comprising more than 40 specialized cell types (Franks T.J. *et al*., 2008, Tata P.R. & Rajagopal J., 2017). The precise orchestration of diverse cell types is indispensable for lung structures and functions (Xu Y. *et al*., 2016). Each of these cell types in a unique signaling environment bears a distinct transcriptome in response to microbial and environmental insults (Cohen M. *et al*., 2018, Reyfman P.A. *et al*., 2019). Dynamic cross-talk between cells and cell-matrix interactions are required in the anatomy integrity and specific functions of the lung (Cohen M. *et al*., 2018, Schiller H.B. *et al*., 2019). The immune cells interacting with tissue components are essential for the lung developmental processes and cell fate decisions (Cohen M. *et al*., 2018, Farber D.L. & Sims P.A., 2019). The interactive gene networks among multiple lung cell types govern the structure and coordinate the cells fate decisions, injury repair, and regeneration (Ardini-Poleske M.E. *et al*., 2017). So, cell-cell communication signals in the lung are intractable to seize because of their high cellular complexity and sensitivity to microenvironments (Amit I. *et al*., 2016, Miller A.J. & Spence J.R., 2017). However, cross-talk of diverse cell types in the lung is currently much less understood. Therefore, cellular communication and genetic regulatory networks (GRNs) in normal physiological and pathological settings remain an active research field of the lung.

Unlike conventional bulk methods, the single-cell RNA sequencing (scRNA-seq) enables the gene expression of an individual cell (Chambers D.C. *et al*., 2019, Saliba A.E. *et al*., 2014), which can reveal cellular compositions and heterogeneity in complicated tissues and molecular pathways specific to a certain cell type (Angerer P. *et al*., 2017, Fu Y. *et al*., 2020, Griffiths J.A. *et al*., 2018, Papalexi E. & Satija R., 2018, Svensson V. *et al*., 2018). The scRNA-seq is especially amenable to reliable identification of novel or rare cell types (Farber D.L. & Sims P.A., 2019) and even closely-related cell populations (Paul F. *et al*., 2015). The unprecedented cellular analysis by scRNA-seq spawns the ambitious international plan called Human Cell Atlas (HCA) (Regev A. *et al*., 2017). As a part of HCA, a comprehensive cell atlas of the human lung in health and disease are currently underway (Chen Z. *et al*., 2020b, Plasschaert L.W. *et al*., 2018, Reyfman P.A. *et al*., 2019, Schiller H.B. *et al*., 2019, Villani A.-C. *et al*., 2017, Xu Y. *et al*., 2016). The unprecedented cellular analysis by scRNA-seq has been recently achieved in multiple tissue and organs (Raredon M.S.B. *et al*., 2019). The cell landscape in mice lung during health and disease are used to elucidate the normal development, homeostasis, and pathogenesis (Angelidis I. *et al*., 2019, Aran D. *et al*., 2019, Dong J. *et al*., 2018, Lee J.H. *et al*., 2017, Montoro D.T. *et al*., 2018, Plasschaert L.W. *et al*., 2018, Treutlein B. *et al*., 2014, Wang Y. *et al*., 2018, Whitsett J.A. *et al*., 2019, Xie T. *et al*., 2018). The rapid increase in the application of scRNA-seq has produced large datasets. The massive published data of human and mouse lung molecular atlas is offered on the LungMAP website, created by the National Heart Lung and Blood Institute, to serve as a research platform and public education tool (Ardini-Poleske M.E. *et al*., 2017). Besides, scRNA-seq data, in combination with computational analyses, can be employed for investigating the cell-cell interactions. The scRNA-seq data was innovatively used to delineate the interactions of immune cells with tissue components in the normal development and fibrotic lung (Farber D.L. & Sims P.A., 2019). A multiplexed network deduced from the adult lung scRNA-seq data was created to capture potential cell-cell interactions (Raredon M.S.B. *et al*., 2019). Using protein-ligand interaction networks, Cohen et al. (Cohen M. *et al*., 2018) analyzed scRNA-seq data to obtain cell-cell interactions in mouse lung development. Base on the scRNA-seq profiles, the interactions between lung adenocarcinoma cells and T cells were explored, and 114 ligand-receptor pairs were identified (Chen Z. *et al*., 2020a).

Respiratory infections are recognized as one of the most severe diseases of growing pigs and lead to massive economic losses (Maes D. *et al*., 2000). Multiple pathogens contribute to a polymicrobial infection (Sunaga F. *et al*., 2020). Significantly, the S protein of severe acute respiratory syndrome coronavirus 2 (SARS-CoV-2) can bind to the cellular receptor angiotensin-converting enzyme II (ACE2) of the pig to mediate viral entry (Liu Y. *et al*., 2020). Although some of the porcine respiratory pathogens are well characterized, the provoked host response and immunological mechanisms are much less understood (Thacker E.L., 2001). Because of the comparable or nearly identical organ sizes and physiology to human beings, pigs have been increasingly considered an optimal source of human organs for transplantation (Sykes M. & Sachs D.H., 2019). The major barrier of porcine xenotransplantation is mainly limited by organ rejection due to incompatibilities between pigs and humans (Shin J.S. *et al*., 2014). By CRISPR-Cas9 and transposon technologies, pigs were genetically engineered to endow the PERVKO·3KO·9TG pigs with greater compatibility with the human immune system (Yue Y. *et al*., 2020). Genetically engineered pig is gradually regarded as an excellent model for studying human diseases, but the translational counterparts in pig suitable for the validated markers for human lung diseases are scarce (Meyerholz D.K. *et al*., 2016). Many scRNA-seq works of human and mouse lungs during normal development or diseases have been reported. Still, there are significant differences between the morphology and cell types during lung development (Ardini-Poleske M.E. *et al*., 2017, Cohen M. *et al*., 2018) and among mammalian species lung (Schiller H.B. *et al*., 2019, Tata P.R. & Rajagopal J., 2017). Therefore, considering the above problems, it is urgent to study cell types, GRNs, cell-cell communications in pig, as well as the conservation and divergence of cell types between pig and human lungs.

Here, we applied scRNA-seq technology to study the cellular heterogeneity of 3 months old domestic pig lung and identify the special marker of each cell type. We constructed cell-specific transcription factor regulatory networks in all cell clusters and further analyzed the cell-cell communications. Importantly, we established an interactive online platform exploring lung single-cell data from four mammalian species (human, mouse, rat, and pig).

## Results

### Construction of domestic pig lung atlas

To efficiently acquire single cells from lung tissues, we used scRNA-seq by the DNBelab C4 to subject lung tissues (Figure 1a). These cells were sequenced by DNBSEQ-G400, and we achieved 13,580 cells. After quality filtering for these cells according to nFeatures and nCount, these cells were performed to 34 clusters by using the gene expression matrix. Next, we annotated the cell types for these clusters based on the markers’ expression in different clusters. In total, we identified 34 clusters corresponding to mainly 16 cell types. Alveolar Epithelial Type 1 (ATI) was characterized by the specific expression of *SAMD12*. Alveolar Epithelial Type 2 (ATII) was indicated by the specific expression of *NAPSA*. Moreover, we obtained Signaling Alveolar Epithelial Type 2 (*TMEM243*), Alveolar Fibroblast (*PTPRD*), Endothelial Cell (*FLT4*), Ciliated Cell (*CFAP44, DNAH5*), Capillary Cell (*CSF1*), Capillary Aerocyte (*TRNP1*), Artery Cell (*ACKR1*), Mucous (*NDNF*) and Secretory Cell (*BPIFA1*) (Figure 1c). In addition to those cells, we also identified several immune cell types. Briefly, Macrophage has a high expression of *MARCO*, and Alveolar Macrophage demonstrated enrichment of *FCER1G*. T cells showed an increased expression of *CD2* and *EPHB6*. B cells were annotated due to the enrichment of *BLINK* and *MEF2C*. Besides, Cell Cycles was indicated by the specific expression of *KIF2C* and *SGO2*. Meanwhile the Uniform Manifold Approximation and Projection (UMAP) plot was used for visualization (Figure 1b). After annotation, we selected the top-10 ranking markers of each cell type for goterm analysis, and the bubble plot was used for goterm visualization (Figure 1d). The result verified our annotation. Of note, most of goterm had a significant correlation with the cell type. For example, cilium development, intraciliary transport, intraciliary transport were involved in the cilium assembly, while the epithelial cilium movement and protein localization to cilium had an obvious significance in the ciliated cell.

**Figure 1:**
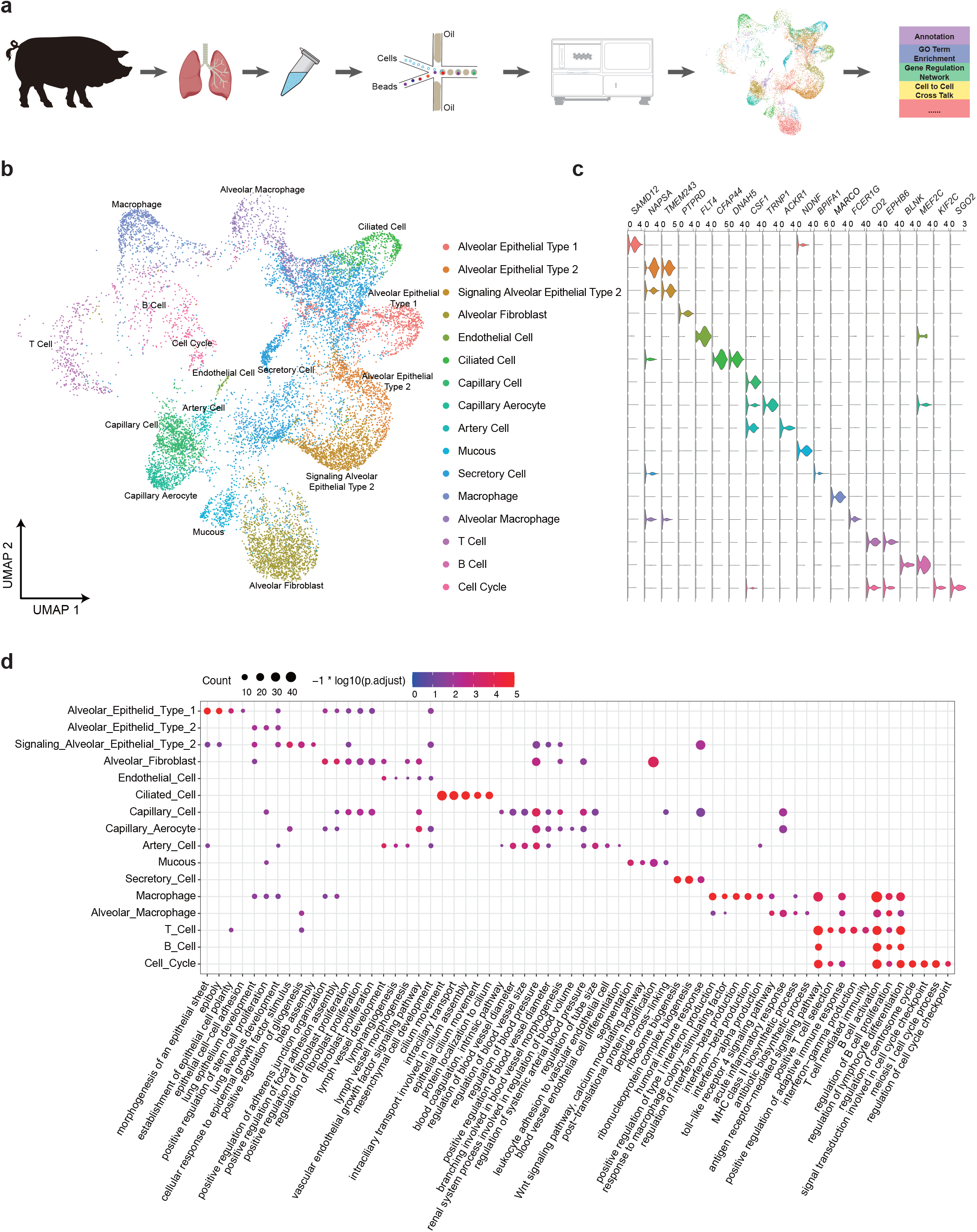
Single-cell atlas of pig lung. (a) Pipeline of article design. (b) UMAP plot of lung cells, color coded for cell types. (c) Violin plots showing the expression of markers, the highest expression indicated the cell types. (d) Bubble plot showing goterms in cell types, color represents the average normalized expression level, and size indicates the percentage of cells.

### Conservation of GRNs between domestic pig and human lungs

By comparing with the single-cell data of human lungs in other articles (Raredon M.S.B. *et al*., 2019), we found that human and domestic pig had conserved transcription factors (TFs) in the main cell types of lungs, including ATI, ATII, Macrophage, Alveolar Macrophage, T Cell, and B Cell. The number of conserved TFs and regulated genes in different cell types between human and domestic pig lungs was shown in Figure 2a-b. Figure 2c shows that ATII has 16 conserved transcription factors (MECOM, EHF, TSC22D3, SREBF2, GRHL2, FOXP2, ETS1, SP110, BHLHE40, NFIX, ETV5, AHR, REL, HMGB2, TCF4, and ZEB2), and they conservatively regulate 2615 target genes. After analyzing the related genes conservatively regulated by the transcription factor ETV5 in ATII, it is found that *MID1IP1* and *NAPSA* genes are specifically and highly expressed in this cell type (Figure 2d). Therefore, we predict that these two genes and transcription factor ETV5 have an essential relationship with the development and function of ATII. The results of the transcription factor regulatory network conserved in other cell types are shown in Figure S2.

**Figure 2:**
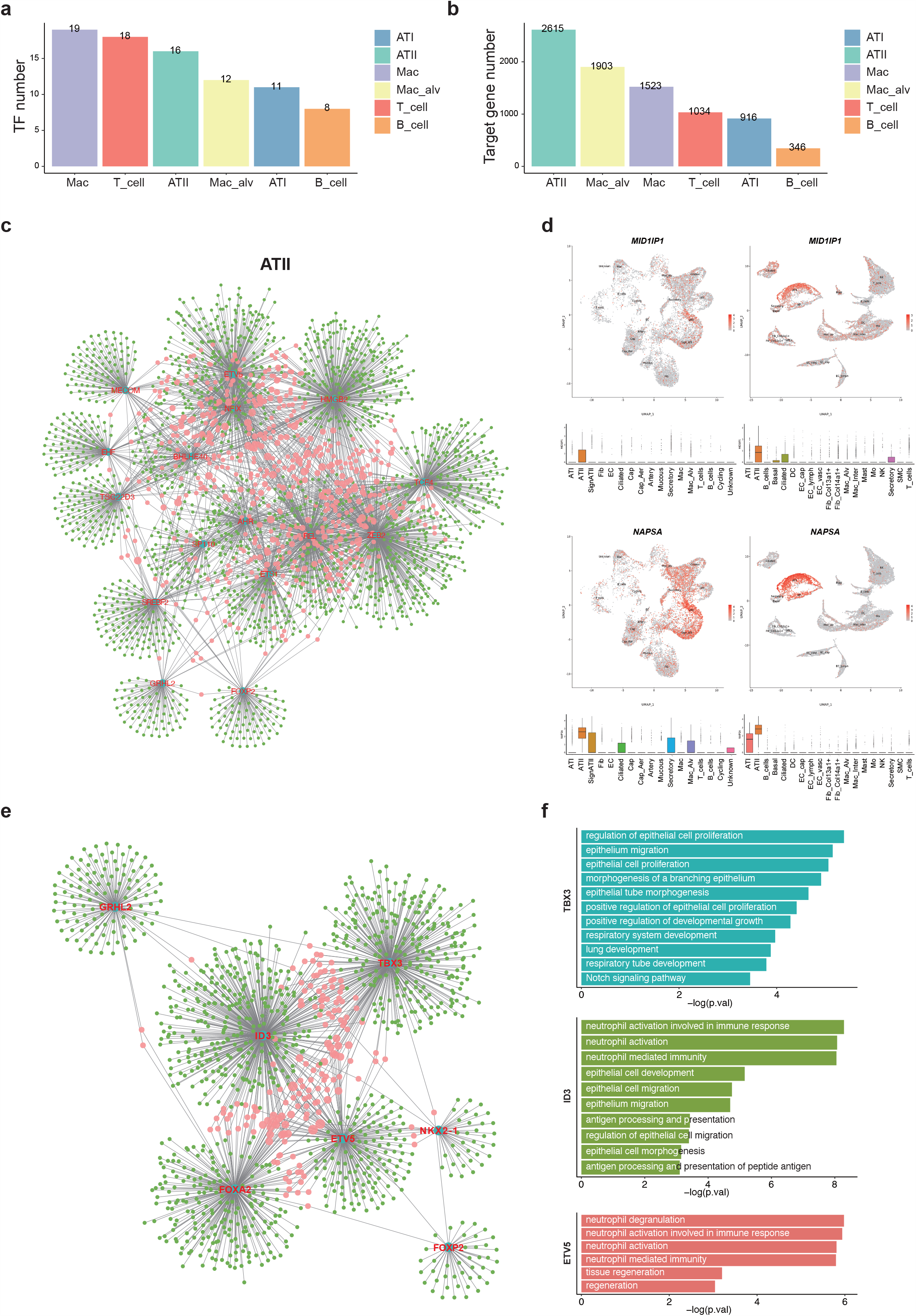
GRNs of TFs in common cell types between pig and human lungs. (a) Number of TF conserved in the lung between pig and human (b) Number of target gene conserved in the lung between pig and human (c) Conservative TF in ATII. Blue nodes indicating TFs, while red and green nodes indicating genes that were considered co-variable with these TFs. Red nodes showed genes that co-variable with more than one TFs, and the size of these red nodes was positive correlated with the number of its co-variable TFs. (d) FeaturePlot of the target gene in ETV5. (e) More TF in ATII. (f) GO terms of TFs co-variable genes of each ATII cell’s development related TFs.

Since our conservative transcription factors were obtained from DEGs, some important transcription factors may be filtered. To make the results more comprehensive, we got the transcription factors (GRHL2, ID3, TBX3, FOXA2, FOXP2, ETV5, and NKX2-1) that are closely related to the development, differentiation, and cell function of human lung ATII from the research articles (Aspal M. & Zemans R.L., 2020, Swarr D.T. *et al*., 2019, Whitsett J.A. *et al*., 2019), and studied the performance of these transcription factors in domestic pig lungs (Figure 2e). We can find ID3 regulates the largest number of genes. Through GO function enrichment analysis, TBX3, ID3, and ETV5 showed functions related to lung development, epithelial cell development, and immune function in domestic pig lung (Figure 2f).

### Comparison of cell-cell communications and intermolecular signaling in domestic pig and human lungs

The communication between cells plays an important role in the structural and functional for maintaining the tissues’ function and development (Pavlicev M. *et al*., 2017). To investigate cell to cell cross-talk relationship of ten main cell types in domestic pig lung, we performed cell communication analysis by using R package CellChat. 600 significant ligand-receptor pairs were detected. And these pairs were further classed into 46 signaling pathways, including VEGF, SEMA3, TGFβ, WNT, IFN-II, IL1, IL4, and IL6, etc. To further understand the roles of all these signaling pathways, we performed global communication patterns recognition analysis (Figure S3a-b) of these signaling pathways by non-negative matrix factorization to identify the global communication patterns and the key signals in different cell groups (Figure 3a).

**Figure 3:**
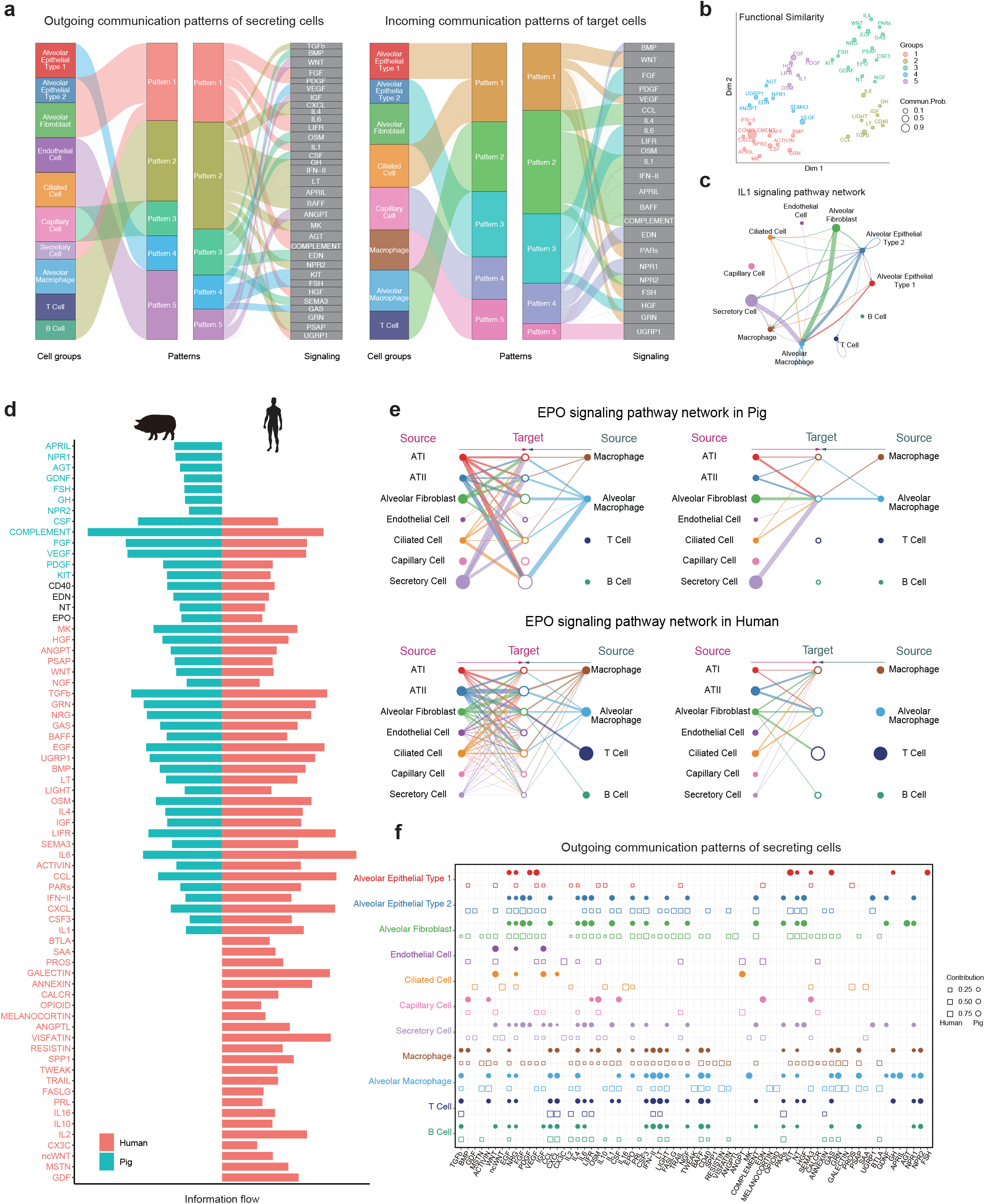
Cell-cell communications in lungs between pig and human by CellChat analysis. (a) The alluvial plot showing outgoing signaling patterns of secreting cells (left plot) and incoming signaling patterns of target cells of pig lung (right plot). (b) Projecting signaling pathways onto a two-dimensions manifold according to their functional similarity. (c) Circle plot showing the inferred IL1 signaling network. (d) All significant signaling pathways were ranked based on their differences in overall information flow within the inferred networks between pig lung and human lung. The top signaling pathways colored blue are more enriched in pig lung, the middle ones colored black are equally enriched in pig lung and human lung, and the bottom ones colored red are more enriched in human lung. (e) Hierarchical plot showing the inferred intercellular communication network of EPO signaling in pig lung and human lung, respectively. The solid indicates the cell type that produces the ligand, the hollow indicates the cell type that produces the receptor. And the direction of the arrow is from source to target. (f) The dot plot showing the comparison of outgoing signaling patterns of secreting cells between pig lung (circle) and human lung (square). The dot size is proportional to the contribution score computed from pattern recognition analysis. A higher contribution score implies the signaling pathway is more enriched in the corresponding cell group.The solid indicates the cell type that produces the ligand; the hollow indicates the cell type that produces the receptor. And the direction of the arrow is from source to target.

And we identified 5 patterns for outgoing patterns and 5 patterns for incoming patterns. The outgoing signals of immune cells were characterized by pattern #2, representing such pathways as IL4, IFN–II, MK, NPR2, GRN, PSAP, etc. (Figure 3a). On the other hand, the communication patterns of target cells (Fig. 3a) shows that incoming immune signaling is dominated by two patterns #2 and #5, which include signaling pathways as CCL, IL4, IL1, IFN–II, APRIL, BAFF, COMPLEMENT, and UGRP1 (Figure 3a). The outgoing ATI signaling was characterized by pattern #4, representing such pathways as VEGF, KIT, FSH, and GAS. ATII, alveolar fibroblast, and secretory cell outgoing signaling were characterized by pattern #1, representing such pathways as FGF, PDGF, IL6, LIFR, IL1, and HGF, etc. The incoming ATI and ciliated cells signaling pathways were characterized by pattern #1, representing such pathways as WNT, VEGF, PARs, FSH, and GRN. The incoming ATII and alveolar fibroblast signaling pathways were characterized by pattern #3, representing such pathways as FGF, PDGF, IL6, LIFR, OSM, and HGF.

Further, the significant signaling pathways were grouped based on their cellular communication network functional similarity. The application of functional similarity grouping identified 5 groups of pathways (Fig. 3b). Group #1, which includes CXCL, GRN, and BMP pathways, largely represents signaling from Secretory cells and Alveolar Macrophage cells. Group #2 is dominated by inflammatory pathways (e.g., TGFβ, CCL, IL6) and largely represents paracrine signaling from Alveolar Macrophage cells to ATI, ATII, and Ciliated cells. Group #3, which includes EPO, IL4, EGF, and WNT pathways, represents signaling from Ciliated cells and Secretory cells autocrine signaling. Group #4, which includes SEMA3, VEGF, NPR1, and EDN pathways, shows a vital role in the lung growth, representing the signaling from ATI, while group #5 includes HGF, PDGF, IL1, and FGF pathways. IL1 pathway exhibited abundant signaling interactions among alveolar macrophage, macrophage, secretory cell, alveolar fibroblast, and ATII. And IL1 ligands were dominantly secreted by secretory sell, alveolar fibroblast, and ATII (Figure 3c).

Moreover, we compared the information flow of different signal pathways between humans and domestic pigs. We found that some signaling pathways were similarly expressed in domestic pigs and humans, including CD40, EDN, NT, and EPO (Figure 3d). The network of CD40, EDN, and NT signaling pathways were shown in Figure S4. From the performance of the EPO signaling pathway network between domestic pigs and humans, it can be found that the trend of this pathway is relatively consistent among different cell types (Figure 3e). We believe that these pathways are equally important in lung development in both species. Compared with domestic pigs, human has inconsistent performance in some signaling pathways: (i) closed (For example, APRIL, NPR1, AGT, etc.), (ii) reduced (such as CSF, FGF, VEGF, etc.), (iii) opened (IL 16, 1L10, IL2, etc.) or (iv) increased (WNT, TGFβ, IL4, etc.). We used pattern recognition analysis to study the detailed changes in the efferent signals of ligand levels in all critical pathways between domestic pig and human and discovered the conservation and difference in signal pathways between them (Figure 3f). NRG, VEGF, SEMA3 in ATI maintain their outgoing signaling patterns, are closely related to lung development. ATII and Alveolar Fibroblast show more conservation signaling pathways between these two species, including EGF, NRG, FGF, PDGF, IL6, LIFR, IL1, EPO, CSF3, LIGHT, NGF, COMPLEMENT, NT, and HGF. When it comes to Macrophage and Alveolar Macrophage, they show a lot of similarities in signal pathways. But there are more differences in T cells and B cells. The pathways maintained in Macrophage and Alveolar Macrophage include TGFβ, ACTIVIN, EGF, IL4, CSF, IFN-II, LT, LIGHT, BAFF, CD40, GAS, GRN, and PSAP. Relatively, T cells of human open IL2 and ANNEXIN signaling pathway. B cells of human open MSTN, IL2, LIFR, IL10, PRL, VISFATIN, ANNEXIN, GRN, and BTLA pathway.

### Construction of the ScdbLung database

To get the utmost out of the lung single-cell resources, we created an online platform named ScdbLung (http://120.79.46.200:81/Lung/index.php). 916558 cells were obtained from 78 datasets for four species (Human, Mouse, Pig, and Rat) (Figure 4a). Cell numbers range from 116 to 114,396. Briefly, 58 datasets, 754766 cells were obtained for human, 15 datasets, and 125807 cells were obtained for mouse, 2 datasets, 10953 cells were collected for rat, 3 datasets, and 25032 cells were obtained for domestic pig. Technology generating those datasets includes 10× Genomics, DNBelab C4, Smart-seq2, Microwell-seq, drop-seq, and inDrop (Table S2).

**Figure 4:**
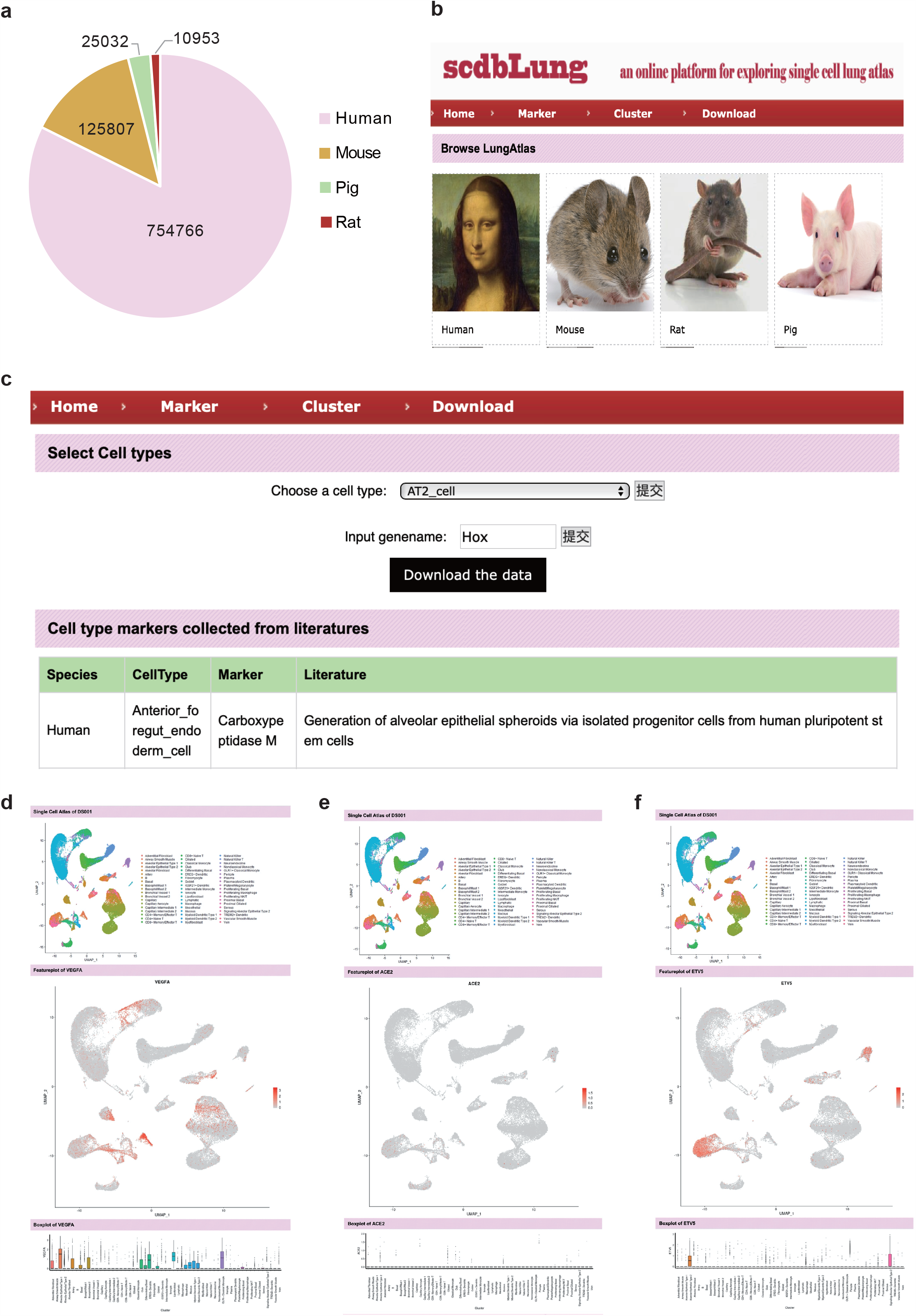
ScdbLung database information and modules. (a) Veen plot showing the percentage of the number of cells in four species (Human, Mouse, Pig, and Rat). (b) The basic framework of ScdbLung (c) The Marker module of ScdbLung. (d) The Cluster module of ScdbLung.

ScdbLung was composed of three functional modules: Marker, Cluster, and Download (Figure 4b). Briefly, the Marker module allows users to explore the marker of different cell types in different species. In total, marker genes for 71 cell types (ATII_cell, ATI_cell, Ciliated_cell, Secretory_cell, B_cell, T_cell, etc.) collected from published literature were included the marker module. Two types of query strategies were provided: firstly, users can search by choosing a specific type, then all collected markers for the selected cell type will be returned. Alternatively, users might be interned in cell types expressing a given marker gene. In that case, we provided a choice for the user to query by gene name, which will return all the cell types, specifically expressing the queried marker (Figure 4c). In the Cluster module, the visualization of cell types of each data is shown, allows users to explore the same gene expression in different data.

Meanwhile, the Cluster module records the data sources and the number of cells (Figure 4d). For example, dataset DS001 displays the human lung atlas constructed using 10× Genomics, comprising 65662 cells from 57 cell populations. Although the expression of cluster-specific genes has been presented in the preprinted article, it isn’t easy to explore the detailed expression patterns of every single gene in each cluster. To solve this problem, we downloaded the matrix data from links provided by the article and plotted boxplot and featureplot for each gene included in the matrix. Subsequently, we presented those plots in the webserver and allowed users to search for any gene they are interested in. For example, if the user selects one dataset, a detail page named “showDS.php” will be returned, mainly composing of two sections. In section 1, the basic information of the dataset will be displayed in the top panel of the webpage, where Species, number_of_cells, Title, Journal, Technology, Data_source, and dataset ID information will be shown so users can fully understand the background of the investigated dataset. In section 2, the UMAP plot showing the distribution of lung cells and cell type labels was provided. A query box was presented to search the expression patterns of any gene by inputting gene symbols. For example, if the keyword “*VEGFA*” was submitted, then the page will jump into the “show.pic” where the boxplot and featureplot of the corresponding gene will be demonstrated. The download module allows users to download the markers and the gene expression matrix of each species. The users can excavate more information and learn by using these data.

## Discussion

The rapid responses to pathogenic assaults and physiological stimuli are attributed to the exquisite coordination of dramatically various cell types in the lung immune system (Sunaga F. *et al*., 2020). In this study, we applied the independently developed sequencing platform by BGI to perform scRNA-seq for 3 months old domestic pig lung, generating the single-cell atlas of 13,580 cells covering 16 major cell types matched with 34 cell clusters. Based on this resource, we systematically characterized the cellular diversity, gene expression heterogeneity, GRNs, and intercellular communication signaling. Recently, the scRNA-seq for the 3.5 months old pig lung was also completed with 11,452 cells by the 10×Genomics platform, which was mainly used to identify the conserved patterns of cell-cell cross-talk in adult mammalian lungs and focus on the roles of alveolar type I cells in tissue homeostasis (Raredon M.S.B. *et al*., 2019). Humans and pigs do not generally cease growing before sexual maturity. The cellular populations and heterogeneity are specific during different stages of development (Szabo P.A. *et al*., 2019). So, our study, together with the 3.5 months old pig lung data (Raredon M.S.B. *et al*., 2019), complementally promoted the comprehension of abundant cell heterogeneity, complicated GRNs, and multicellular interplay in the developmental pig lung.

GRNs of cell type-specific TFs and their candidate target genes might provide an essential basis for cell functions. Here, we predicted the conserved TFs and their candidate target genes in each of the basic cell types between domestic pig and human lungs and found that ATII cells held the most candidate target genes (Figure 2b). ATII cells proliferate to create new ATII cells and differentiate into ATI cells for alveolar regeneration during injury, critical to the stereotyped gas exchange function and normal barrier (Aspal M. & Zemans R.L., 2020). So, we chose ATII cells as the representative for analyzing the conserved GRNs across two species. Both 16 common TFs and 2615 candidate target genes, expressed explicitly in ATII cells of human and pig, were exquisitely orchestrated into the intricate regulatory networks. Noticeably, *NAPSA*, and *MID1IP1* genes as the targets of ETV5 are specifically and highly expressed in ATII cells. NAPSA protein was expressed and secreted from ATII cells in human (Lindskog C. *et al*., 2014). ETV5 in ATII cells was implicated in ATII-to-ATI cell differentiation during lung regeneration (Zhang Z. *et al*., 2017), which was in accord with our GO function enrichment analysis. The highly conserved GRNs in domestic pig and human lungs indicates that extremely large amounts of data on human lung provides guidance for pig research. In general, the GRNs identified in this study could enhance our understanding of gene expression and regulation in ATII cells, which provides a method for GRNs in other cell types.

The complex cell-cell commnnication and intermolecular signaling, coordinated by diverse cell types, were involved with the maintenance of lung-specific structure and functions during physiological and pathogenic processes (Farber D.L. & Sims P.A., 2019, Puttur F. *et al*., 2019) and against environmental and microbial insults (Schiller H.B. *et al*., 2019, Whitsett J.A. *et al*., 2019). Analysis of scRNA-seq data using protein-ligand interaction networks profiled the complex cell-cell interactions during normal lung development of mice (Cohen M. *et al*., 2018). Base on all known ligand-receptor signaling, scRNA-seq data of the adult lung across four species were fully harnessed to delineate the potential conserved intercellular interactions network (Raredon M.S.B. *et al*., 2019). In patients with COVID-19, the severe cases exhibited strong interactions between epithelial and immune cells. The expression of the SARS-CoV-2 entry receptor ACE2 in epithelial cells of patients increased on average three-fold, which correlated with interferon signals by immune cells (Chua R.L. *et al*., 2020). Here, we leveraged our scRNA-seq data of domestic pig lung to find that the IFN-II signaling pathways in macrophage and T cell were similar between domestic pigs and humans (Figure 3f, Figure S4d). Therefore, we inferred that the conserved IFN-II signaling pathway might be implicated in the SARS-CoV-2 entry into human and pig, which was in good accordance with the ability of pig ACE2 to support SARS-CoV-2 entry *in vitro* (Liu Y. *et al*., 2020). In addition, we identified the intercellular cross-talk of 10 major cell types in domestic pig lung. We found some similarities and differences in signaling pathways and their efferent signals of ligand levels between domestic pigs and humans. The interplay among immune cells from our cell-cell cross-talk network would provide clues to eliminate major xenoantigens and enhance the immunological compatibility between pigs and humans for improving porcine xenotransplantation with the next generation of engineered 3KO·9TG pigs (Meyerholz D.K. *et al*., 2016, Yue Y. *et al*., 2020).

The HCA consortium aims to profile all single cells in the human tissues, and lung tissue has priority in the ambitious projects (Schiller H.B. *et al*., 2019). The Human Lung Cell Atlas has been contributed by several programs (Schiller H.B. *et al*., 2019), such as LungMAP in the healthy lung. The LungMAP website (www.lungmap.net), including lung images, transcriptomic, proteomic, and lipidomic data of human and mouse, has been openly available since early Fall of 2017, which will be useful in the understanding of lung development (Ardini-Poleske M.E. *et al*., 2017). Recently, the scRNA-seq has allowed unprecedented advances in dissecting cellular heterogeneity and transition during lung organogenesis (Dong J. *et al*., 2018, Han X. *et al*., 2018, Lee J.H. *et al*., 2017, Wang Y. *et al*., 2018, Whitsett J.A. *et al*., 2019). This technology is also beneficial in defining cellular alterations in disease models, such as multiple mesenchymal cell types (Xie T. *et al*., 2018, Zepp J.A. *et al*., 2017), alveolar macrophage phenotypes in the fibrotic lung (Aran D. *et al*., 2019, Reyfman P.A. *et al*., 2019), and epithelial cell diversity in idiopathic pulmonary fibrosis (Xu Y. *et al*., 2016). Except lung cellular complexity of human and mouse, Raredon, *et al*. (Raredon M.S.B. *et al*., 2019) created lung atlas from 9 weeks old rat and 3.5 months old pig. So far, a rapid growing single-cell data under physiological and pathological conditions has been generated by researchers around the world. However, a platform dedicated to analyze and explore the cross-species lung single-cell atlas is nearly missing. Here, we presented a comprehensive and freely accessible online database, ScdbLung, which integrated a total of 916558 cells, composing 78 datasets sets from four mammalian species. Our platform will be beneficial to fully exploit the valuable resources of lung scRNA-seq atlas.

There are some apparent disadvantages to the current scRNA-seq techniqu e. This technique loses spatial location information, just like it in our study. Samples collected in different anatomical regions for sequencing and immunostaining information of marker genes have been attempted to deal with this limitation. Tissue sources from different anatomical regions were subjected to scRNA-seq and then discovered differences in lung structure and cell populations among upper and lower airways and alveoli parenchyma (Vieira Braga F.A. *et al*., 2019). According to physical sites of cell type-specific markers from histologic data of some public databases and immunostaining results□two distinct endothelium cell types from scRNA-seq data can be assigned to the histologic location in the lung (Raredon M.S.B. *et al*., 2019). Besides, the scRNA-seq can provide transcriptional information of complex tissues at the single-cell level, which is limited in solving lung development or complex lung diseases. The comprehensive analysis by multi-omics (such as single-cell epigenome, transcriptome, proteome, metabolome, etc.), together with function experiments, are warranted to facilitate the solution of these problems. For example, the scRNA-seq combined with the scATAC-seq (Buenrostro J.D. *et al*., 2018) or mass spectrometry-based proteomics (Angelidis I. *et al*., 2019) can build a more accurate cell-type-specific regulatory network. Base on the massive single-cell sequencing data of lung from several species, innovative computational methods should be customized to resolve a specific scientific problem, such as dissecting cell-cell communication and signaling pathways during lung development and physiological status (Farber D.L. & Sims P.A., 2019). Finally, the lung is constantly exposed to environmental insults and need dynamic cell activities for maintaining pulmonary homeostasis (Farber D.L. & Sims P.A., 2019). Mammalian tissues or organs usually do not stop developing before sexual maturity (Raredon M.S.B. *et al*., 2019). The human lung generally keeps growing after birth to adolescence and beyond (Schiller H.B. *et al*., 2019). There were significant differences in lung anatomy and cell types between mice and humans (Tata P.R. & Rajagopal J., 2017). Therefore, to comprehensively understand the cellular basis of pig lung functions, it is necessary to sample pig lung tissues at different developmental stages or physiological states for scRNA-seq.

## Material and methods

### Ethics statement

Research and sample collection were approved by the Institutional Review Board on Ethics Committee of BGI [Approval letter reference number BGI-NO. BGI-IRB A20008]. All procedures were conducted according to the guidelines of the Institutional Review Board on the Ethic Committee of BGI. All applicable institutional and/or national guidelines for the use and care of animals were followed.

### Sample collection and nuclei extraction

Duroc□×□(Landrace × Yorkshire) 3-way cross pigs (DLY) were purchased from an agricultural market. This research dissected 8 lung tissue pieces from three different 3-month-old male DLY (The sex of the pig did not influence this study). The collected tissues were washed by 1× PBS, then quickly frozen and stored in liquid nitrogen. Nuclei extraction was separated by mechanical extraction method (Liu C. *et al*., 2019). Firstly, put the tissues into 2ml Dounce homogenizer set and thawed in homogenization buffer (containing 20mM Tris pH8.0, 500mM sucrose, 0.1% NP-40, 0.2U/μL RNase inhibitor, 1× protease inhibitor cocktail, 1% bovine serum albumin (BSA), and 0.1mM DTT. Use Dounce pestle A to grind the tissue 10 times, filter with 70μm cell filter, and then grind with Dounce pestle B 10 times, filter with 30um cell filter. Centrifuge at 500 × g for 5 minutes at 4°C to pellet the nuclei, and resuspend in the blocking buffer containing 1% BSA and 0.2U/μL RNase inhibitor in 1× PBS. Centrifuge again at 500 × g for 5 minutes and resuspend with Cell Resuspension Buffer (MGI).

### Single nuclei library construction and sequencing

The mRNA capture was operated on DNBelab C4 device (MGI). Complete cDNA amplification and library construction according to the MGI DNBelab C series reagent kit (MGI). All libraries were further prepared based on the DNBSEQ-G400 sequencing platform. Sample information and quality control are shown in Figure S1 and Table S1.

### Single-cell RNA sequencing data processing

Sequencing data filtered and gene expression matrix was obtained using DNBelab C Series scRNA-analysis-software (https://github.com/MGI-tech-bioinformatics/DNBelab_C_Series_scRNA-analysis-software). The single-cell analysis was conducted using the Seurat v3 (Stuart T. *et al*., 2019). A series of preprocessing procedures were performed separately on each sequencing library before clustering. Normalization and definition of highly variable genes (HVGs) were performed for each sequencing library with default parameters. “FindIntegrationAnchors” and “IntegrateData” functions were applied to integrate all 8 sequencing libraries and reduce the batch effect. Then we integrated our datasets with previously reported pig scRNA-seq datasets (Raredon M.S.B. *et al*., 2019) to facilitate and verify the cell types identification. Principal component analysis (PCA) was performed using HVGs, and principal components (PCs) significance was calculated using the “JackStraw” function. We chose PCs with p.values< 0.01 for downstream cluster identification and visualization. UMAP was applied for visualization.

### Differentially analysis and functional enrichment analysis

Differentially expressed genes (DEGs) were identified using the “FindAllMarkers” function implemented in Seurat84. Wilcoxon rank sum test was applied. Gene with adjusted P-value (Bonferroni method) lower than 0.05 was defined as DEGs.

### Cell type annotation

Cell types were assigned by the expression of known cell-type markers retrieved from published researches.

### Transcription factor regulatory network in domestic pig lung

To investigate the transcription factors regulatory network of the pig lung ATI and ATII cell types, we performed a correlation analysis of 7 known TFs, which are reported important to lung development. The correlation analysis was performed using the corr.test() function of the psych R package. We used p_value >= 0.001 and r >= 0 to get the putative coexpression gene lists of each TFs. Then, we graphed the network by using cytosacape (Aspal M. & Zemans R.L., 2020, Swarr D.T. *et al*., 2019, Whitsett J.A. *et al*., 2019).

### Comparison of transcription factor regulation of main cell types of lung between pig and human

To compare the transcription factor regulation of the main cell types of lung between pig and human, we first get the TFs lists of pig and human from AnimalTFDB 3.0 (Hu H. *et al*., 2019). And a previously reported human lung scRNA-seq dataset (Raredon M.S.B. *et al*., 2019) was introduced for the following comparison. Then we perform correlation analysis (mentioned above) of TFs identified in the differential expression gene list of each cell type among two datasets for six main cell types (including ATI, ATII, B Cell, T Cell, Macrophages, and Alveolar Macrophage). We used the p_value>= 0.05 and r >= 0 to generate the putative coexpression gene lists for each cell type of two datasets. Then a shared putative coexpression gene list was generated for each cell type of pig and human.

### Cell communication and signaling pathways

Cell communication analysis was performed by using R package CellChat (Jin S. *et al*., 2020) with default parameters. Pig and human lung datasets were analyzed separately. Since the absence of pathways database corresponding to the pig for this package, CellChatDB.human was used for both pig and human lung datasets.

### Data Collection and Metadata Annotations

We searched the National Center for Biotechnology Information (NCBI) and The European Bioinformatics Institute (EBI) databases using the following keywords: lung, single-cell, RNA-seq. These data must contain gene expression profiling. The metadata for these cells specifying which cell type each cell barcode belongs to is optional. Meanwhile, we also acquired data from ourselves in CNGBdb. In this study, we had collected 916,558 single cells from 78 datasets (Supplemental Table S2).

Data that provided gene expression profiling but no metadata were processed using the package of Seurat in R. After creating the object, annotated the cell types of each cell using scCATCH package of R or markers which came from related articles or databases.

### Data Visualization

The gene expression profiling and metadata were created to object using the CreateSeuratObject function. Before created these objects, we transformed genes of 11 species into homologous genes in the mouse. The homologous gene correspondence tables were downloaded in BioMart. And we used the Dimplot function to generate the Umap plot with datasets by our cell types. Meanwhile, we used the FeaturePlot function to visualize each symbol’s expression in all cells. These functions are in the Seurat package of R. Finally, the symbol’s expression in every cell type was visualized by geom_boxplot function by the ggplot2 package in R.

## Author contributions

DS.C. and LJ.Z. conceived the idea. LJ.Z. designed and conducted the experiments. JC.Z. and WY.W. assisted with the experiment. JC.Z. analyzed the data. LJ.Z., LH.L., XL.Z. assisted data analysis. HY.W. collected the datasets and analyzed the data of datasets. DS.C. constructed the online database. LJ.Z., JC.Z., HY.W., and XL.Z. wrote the manuscript. DS.C. and XL.Z. revised the manuscript. J.X., P.L., F.C., H.J., and XL.Z. provided a necessary resource for this study. All authors reviewed and approved the final manuscript.

## Acknowledgments

This work was supported by China National GeneBank (CNGB). Our project was financially supported by funding from the National Natural Science Foundation of China (31670742).

## Data Availability

The data have been deposited on the website (http://120.79.46.200:81/Lung/index.php). The data also available in the CNGB Nucleotide Sequence Archive (CNSA: https://db.cngb.org/cnsa; accession number CNP0001486).

---

**Figure S1:**
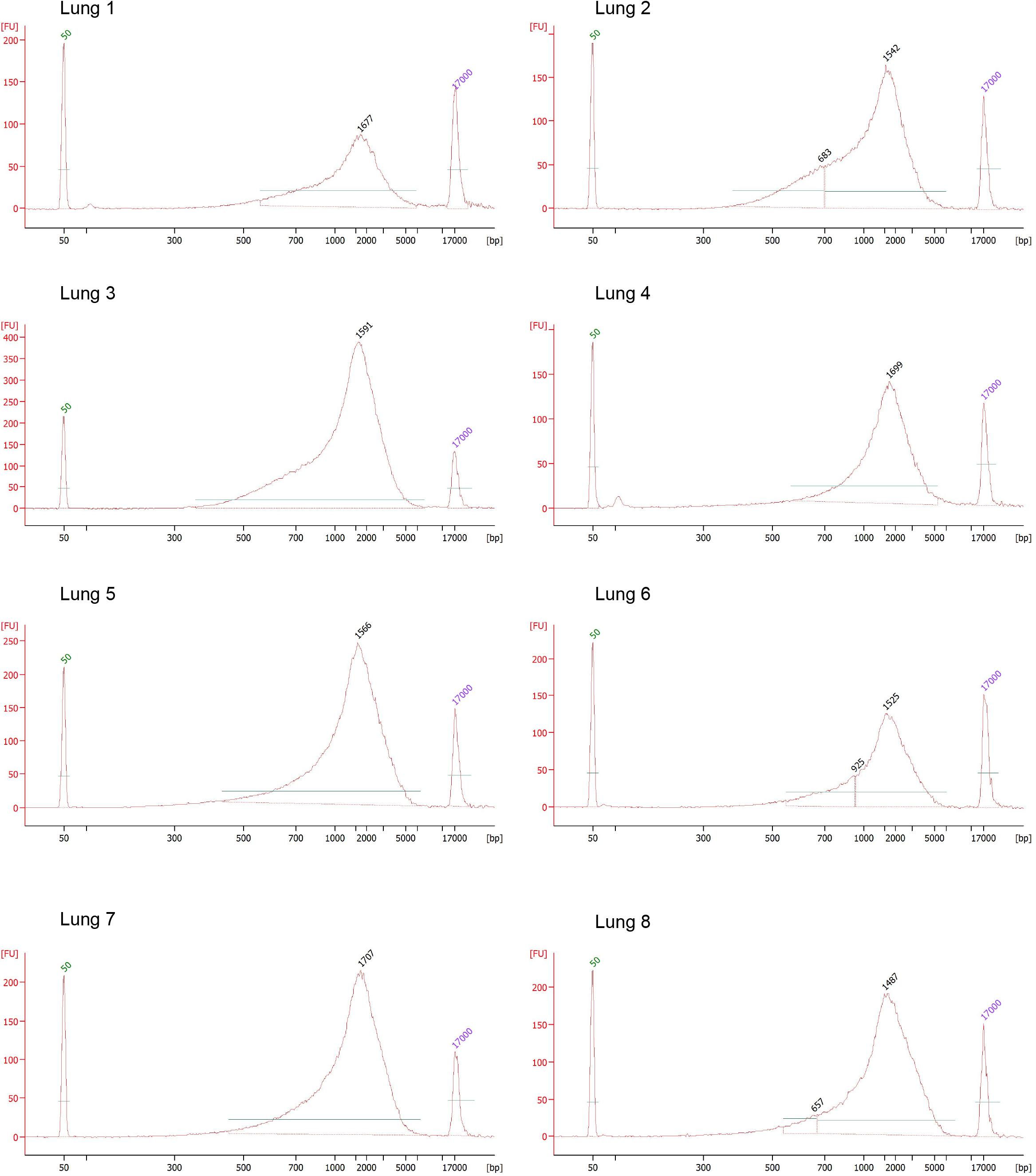
Distribution of cDNA fragment. The visualization of quality control (QC) of lung samples cDNA fragment distribution by using 2100.

**Figure S2:**
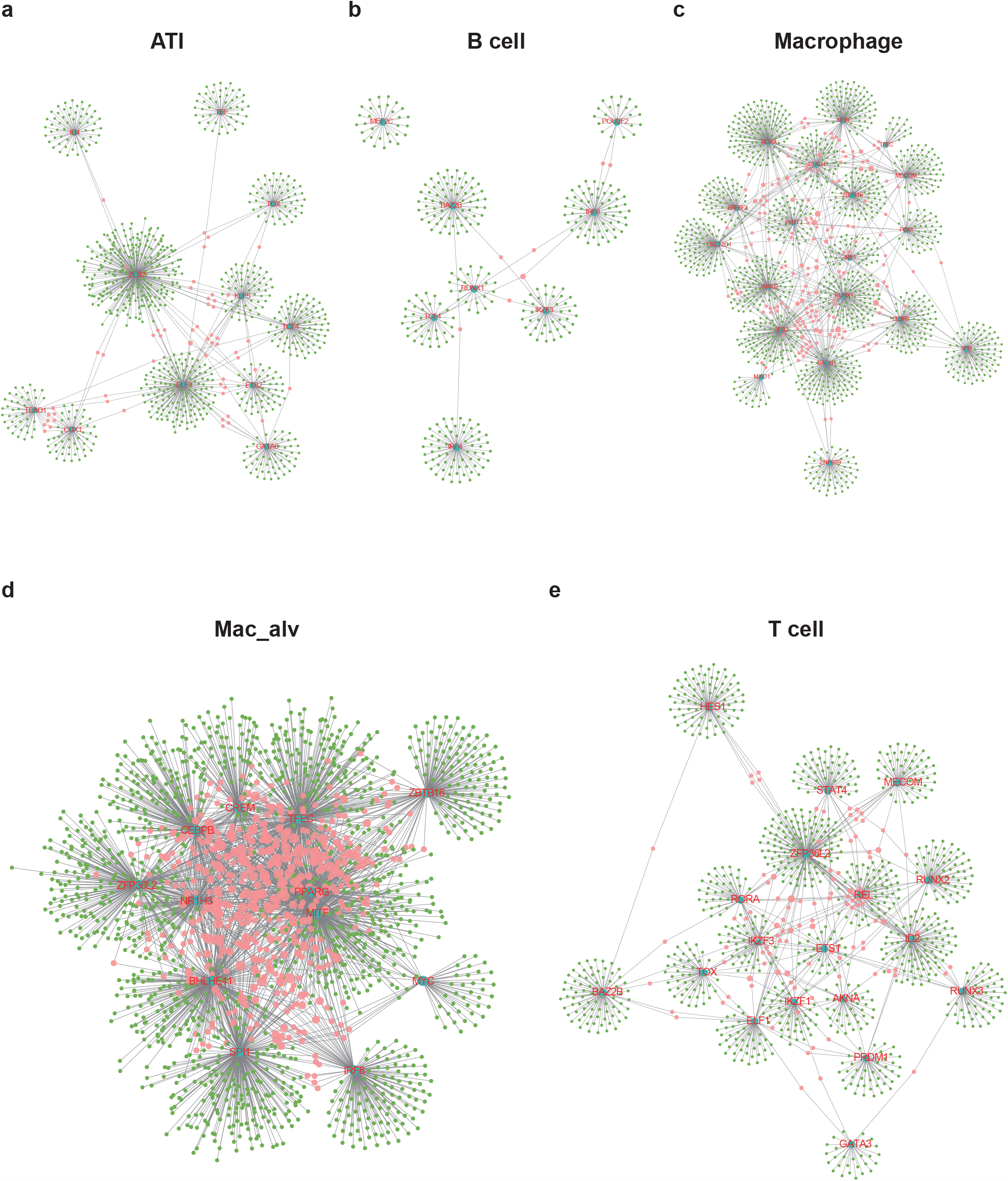
Genetic regulatory network of TFs. Conservative TFs in ATI, B cell, Macrophage, Mac_alv, and T cell.

**Figure S3:**
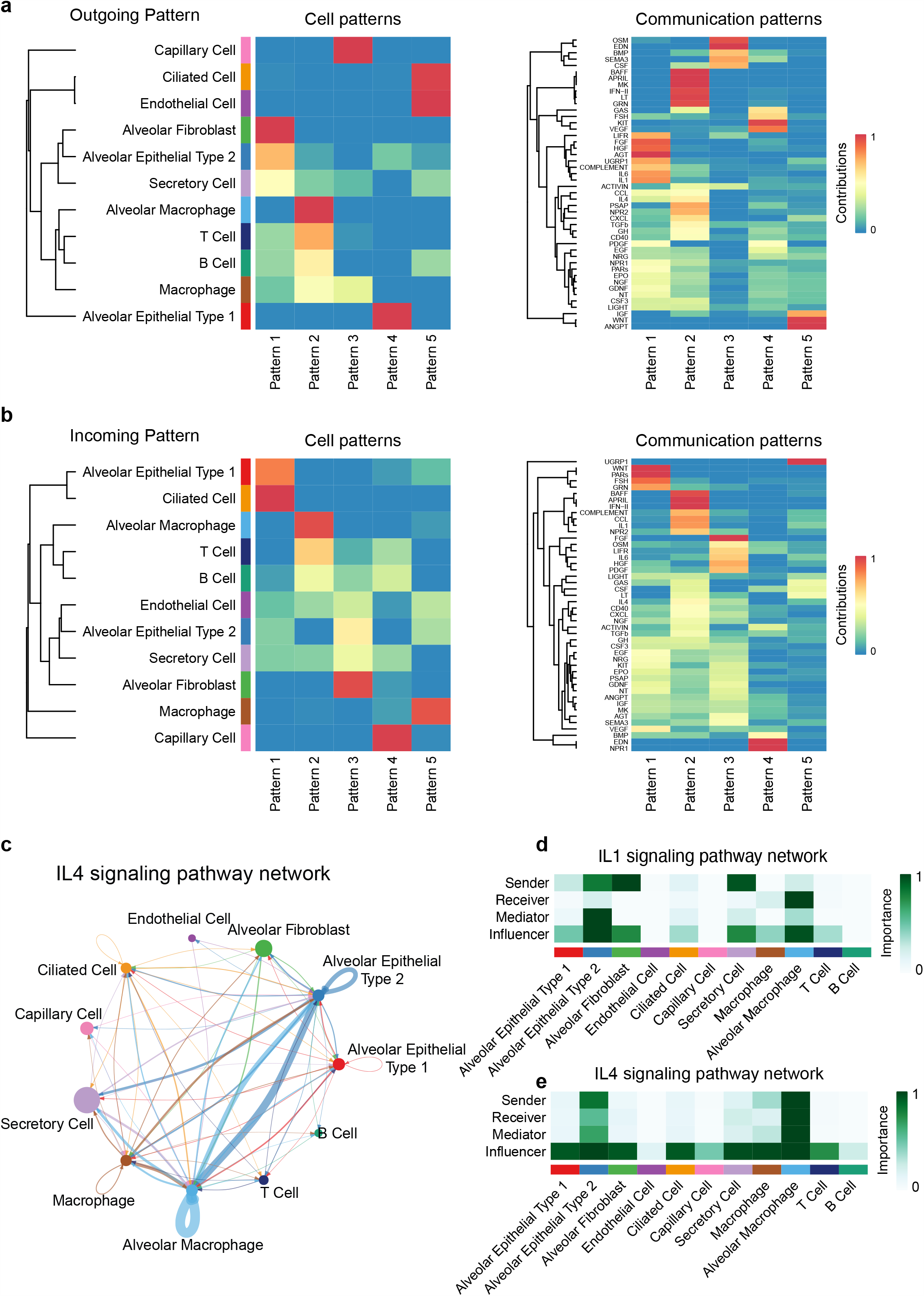
Pattern information and plot of IL4, IL1 signaling pathway networks. (a) Heatmap showing the outgoing pattern among patterns based on cell types and signaling pathways. (b) Heatmap showing the incoming pattern among patterns based on cell types and signaling pathways. (c) Circle plot showing the IL4 signaling pathway network. (d) Heatmap showing the relative importance of each cell type in four networks of the IL4 signaling pathway. The color indicates higher importance in the corresponding network. (e) Heatmap showing the relative importance of each cell type in four networks of the VEGF signaling pathway.

**Figure S4:**
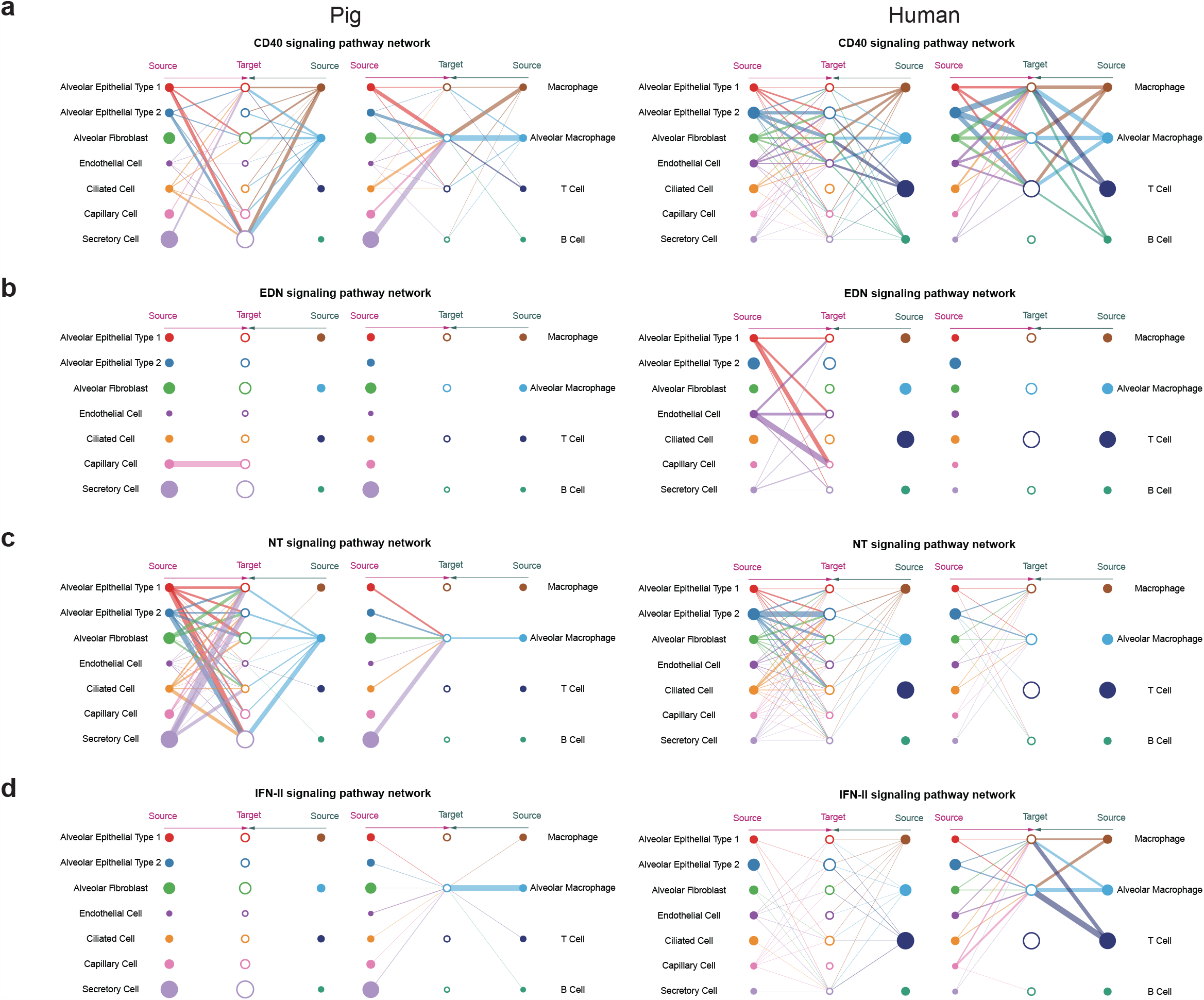
Plots of CD40, EDN, NT, and IFN-II signaling pathway. Hierarchical plot showing the inferred intercellular communication network of CD40, EDN, NT, IFN-II signaling pathway in pig and human lungs.

